# Increased academic performance and prolonged career duration among principal investigators in ecology and evolutionary biology in Taiwan

**DOI:** 10.1101/2022.01.31.478501

**Authors:** Gen-Chang Hsu, Wei-Jiun Lin, Syuan-Jyun Sun

**Author notes:** Corresponding author: Syuan-Jyun Sun.

## Abstract

Academic job markets have become increasingly challenging worldwide, yet it remains poorly characterized how competitively-successful candidates should be and what the underlying determinants of their success are. Focusing on the field of ecology and evolutionary biology, we analyzed the academic performance (measured as h-index) as well as the duration before recruitment as a new faculty member and promotion to full professor of 145 principal investigators (PI) over the past 34 years in Taiwan. We found that PIs had higher performance and longer duration before recruitment more recently. Performance before promotion remained stable, whereas the duration increased over time. The origin and prestige of doctorate had no effect on the performance or duration either before recruitment or before promotion. We also found that the difference in performance before and after recruitment (“After” performance — “Before” performance) decreased in recent years, with PIs recruited in earlier years maintaining their performance after recruitment while those recruited in later years exhibiting a performance drop. While PIs performed equally well before and after recruitment irrespective of doctorate origin, those with domestic PhD degrees showed a decrease in performance after promotion compared to their counterparts with foreign degrees. Taken together, our findings reveal a prolonged career duration for researchers as a result of intensifying competition in academia, and highlight the increasingly crucial role of academic performance, rather than PhD degree itself, in determining academic success.

## Introduction

The academic job market has been increasingly competitive in many fields of science, technology, engineering, and mathematics (STEM) (Cyranoski et al. 2011; Ghaffarzadegan et al. 2015; Xue and Larson 2015), with more PhDs produced but vacancies for tenure-track academic positions remaining relatively constant over the past four decades (Larson et al. 2014; Schillebeeckx et al. 2013). For example, in the US, only 7.6% of new PhDs in life sciences landed tenure-track positions within three years after graduation in 2010. Such a surplus of PhD supply has also emerged in other STEM fields ([NSF] National Science Foundation 2018).

The intensifying competition for tenure-track positions, due to disproportionately high numbers of applicants per position (Larson et al. 2014), has resulted in higher expectations for academic performance shaped by a *“publish or perish”* culture (Garfield 1996). A survey of evolutionary biologists recruited as junior researchers at the National Centre for Scientific Research (CNRS) in France showed that academics recruited in 2013 published nearly twice as many papers as those recruited in 2005 did (Brischoux and Angelier 2015). Additionally, although the minimum education requirement for a tenure-track position is having a PhD degree, it has become increasingly frequent for applicants to have one or even more postdoctoral appointments. Consequently, many PhDs in STEM work as postdoctoral researchers for a prolonged period of time and wait for future opportunities until they are competitive enough in the academic job market (Swihart et al. 2016), whereas some turn to alternative careers outside academia. In the aforementioned CNRS example, Brischoux and Angelier (2015) also found that the time between first publication and recruitment had increased from 3.25 to 8.0 years. The increase in postdoctoral training time can be detrimental to not only the scientific community but also individuals because this increases the age at which researchers become independent, and they have to trade off families for research, with fixed-term and relatively low-paying jobs (Acton et al. 2019).

Despite widely claimed that publication expectations and career duration have surged, empirical quantification of the determinants regarding the change in academic profiles over time remains understudied. In addition to research productivity, which directly predicts the success of recruitment (van Dijk et al. 2014), the origin and prestige of doctoral-granting institutes are critical indicators for academic employment as well (van Dijk et al. 2014), especially in East Asian countries (Shin and Kehm 2013). With the initiative to build world-class universities, many East Asian universities preferentially recruit returnees who obtained PhD degrees from top-ranking universities in Western countries. Hence, competition for limited tenure-track positions is exacerbated when foreign PhDs are favored, leaving domestically-trained PhDs deprived of career development opportunities (Chen 2021). Yet, whether and to what extent publication expectations and career duration differ between domestic and foreign PhDs, and if their academic productivities vary between pre- and post-employment, remain largely unexplored.

In this study, we examined how academic performance as well as duration before recruitment as a new principal investigator (PI) and promotion to full professor changed over time, and how PhD university origin, PhD university ranking, and gender affected the career success. Specifically, we tested the following questions: (1) Is the academic performance for recruitment or promotion associated with the year of recruitment, PhD university origin, ranking, and gender? (2) Is the duration before recruitment or promotion affected by the year of recruitment, academic performance, PhD university origin, ranking, and gender? (3) Does the academic performance of PIs differ before and after recruitment or promotion? To address these questions, we analyzed the data on 145 faculty members in the field of ecology and evolutionary biology in Taiwan between 1987 and 2021. We aim to provide empirical evidence to illustrate the temporal variations in researchers’ publication performance necessary to secure a faculty position and get a promotion, the role of PhD university and gender in determining the success of academic employment, and how these factors contribute to PIs’ future academic performance.

## Materials and Methods

### Data collection

Between November and December, 2021, we surveyed tenure-track faculty members at seven universities in Taiwan, all of which were qualified as research-intensive universities and ranked top 150 in Asia according to 2022 QS Asia University Rankings (https://www.topuniversities.com/). We also surveyed academics from Academia Sinica, a leading academic institution in Taiwan. Together, these eight institutes encompassed 34 academic departments/divisions that serve as tenure homes to the field of ecology and evolutionary biology (including ecology, evolution, biodiversity; see Appendix S1 for details). We excluded researchers in biomedical sciences because publication rates, performance, and collaboration opportunities can vary considerably among these fields (Laurance et al. 2013). A total of 145 PIs who had an updated curriculum vitae online (e.g., institutional/personal websites or Open Researcher and Contributor ID [ORCID]) were identified in our survey, with key information on the university and year of PhD completion, the year of recruitment as a new PI, the year of promotion to full professor, and gender, which is well-documented as a key determinant of performance (Witteman et al. 2019). The university ranking was determined based on 2022 QS World University Rankings. The duration before recruitment as a new PI was calculated as the time between PhD completion and landing a faculty position; the duration before promotion to full professor was calculated as the time between landing a position and getting a promotion.

### Measurement of academic performance

We collected data on academic performance, measured as h-index (Hirsch 2005), from the Publish or Perish software using Google Scholar data, which are freely available and more transparent for tenure reviews (Pauly and Stergiou 2005). We included peer-reviewed papers and book chapters regardless of authorship for calculation of h-index, while PhD theses and conference presentations were excluded. Although other matrices, such as the number of publications and citations, are also commonly used for measuring academic performance, they were both highly correlated with h-index in our study (publications: *r* = 0.91, *p* < 0.001; citations: *r* = 0.77, *p* < 0.001), which had also been found in previous studies (Laurance et al. 2013; Ryan Haley 2012). We thus focused on h-index, a widely accepted measure of academic success that incorporates the assessment of quantity (number of papers) and quality (citations) of publications (Glänzel 2006).

We calculated h-index within the five-year interval both before and after the year of recruitment and promotion, generating up to four h-indexes for each PI. We used the duration of five years because this time span is commonly used by institutes to evaluate the most recent academic performance both for recruiting a new PI and for promotion to full professor. The publications and citations during the year of recruitment and promotion were considered as the performance before recruitment and promotion because these publications, either as published papers or manuscripts “accepted” or “in press”, would most likely contribute to the evaluation of academic performance prior to successful recruitment and promotion. For example, a PI who started a position in 2010 would have an h-index measured for publications between 2006 and 2010 (i.e., “Before” h-index for recruitment), and another h-index measured for publications between 2011 and 2015 (i.e., “After” h-index for recruitment). We did not include “After” h-indexes for PIs who were recruited or promoted less than five years so that all performances have comparable duration.

### Statistical analyses

(1) Academic performance before recruitment/promotion. To examine how various factors affected the academic performance before recruitment as a new PI and promotion to full professor, we fit linear mixed-effects models (LMMs) with PhD university origin (binary variable: Taiwan vs. Foreign), PhD university ranking, year of recruitment/promotion, gender, and all single-factor interactions with year as fixed effects, the institute (department) nested within university as random effects, and the “Before” h-index for recruitment/promotion as the response.

(2) Duration before recruitment/promotion. To examine how various factors affect the duration before recruitment and promotion, we fit LMMs with PhD university origin, PhD university ranking, year of recruitment/promotion, gender, the “Before” h-index for recruitment/promotion, and all single-factor interactions with year as fixed effects, the institute (department) nested within university as random effects, and the duration before recruitment/promotion as the response.

(3) Difference in academic performance before and after recruitment/promotion. To compare the academic performance before and after recruitment and promotion, we fit LMMs with PhD university origin, PhD university ranking, year of recruitment/promotion, gender, and all single-factor interactions with year as fixed effects, the institute (department) nested within university as random effects, and the difference between “After” and “Before” h-index for recruitment/promotion (i.e., “After” h-index — “Before” h-index) as the response.

LMMs were performed using the package “lme4” (Bates et al. 2015); post-hoc pairwise comparisons were performed using the package “emmeans” (Lenth 2021). Response variables (h-index and duration before recruitment/promotion) were log-transformed prior to analyses to meet the assumption of normality. The assumption of independence and equal variance were both assessed using the residual plots. Non-significant interactions (*p* > 0.05) were dropped from our final model results. All analyses were performed in R version 4.1.2 (R Development Core Team 2014).

## Results

In total, we collected data on 145 tenure-track faculty members, of which 44.8% were full professors, 24.8% were associate professors, and 30.3% were assistant professors. Nearly half of the PIs obtained their PhD degrees from the USA (45.5%), followed by Taiwan (33.1%), and relatively few from the UK (4.8%) and other countries (Fig. 1). The PhD universities varied widely in the ranking of prestige among 73 universities from 16 countries (Fig. 2). The gender difference was substantial, with males (112) being around four times as many as females (33).

**Fig. 1.**
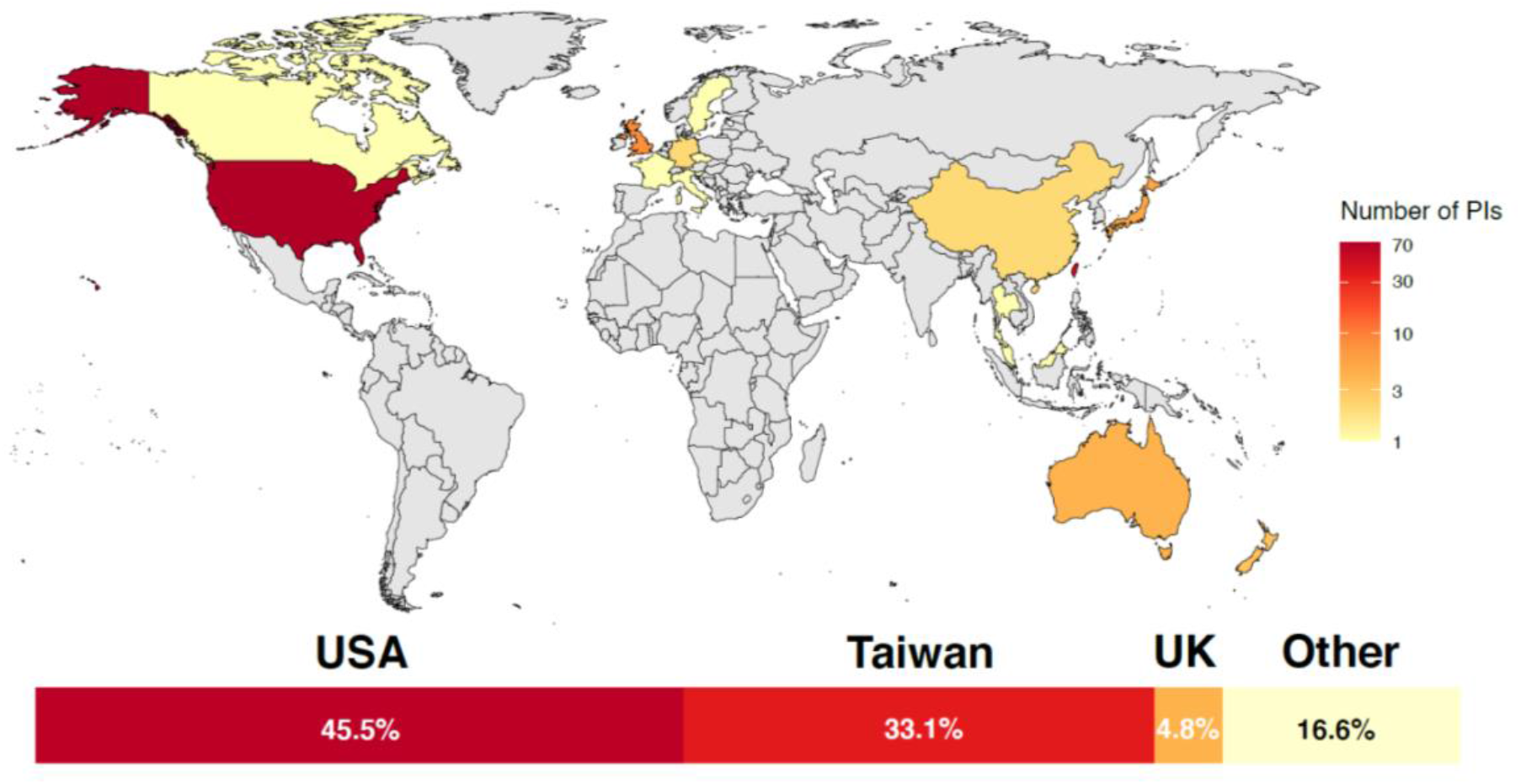
Distribution of the universities from which the 145 PIs obtained their PhD degrees. Percentages of PhD degrees obtained from the USA, Taiwan, and the UK are as noted; “Other” includes all other countries with percentages less than 4.0%

**Fig. 2.**
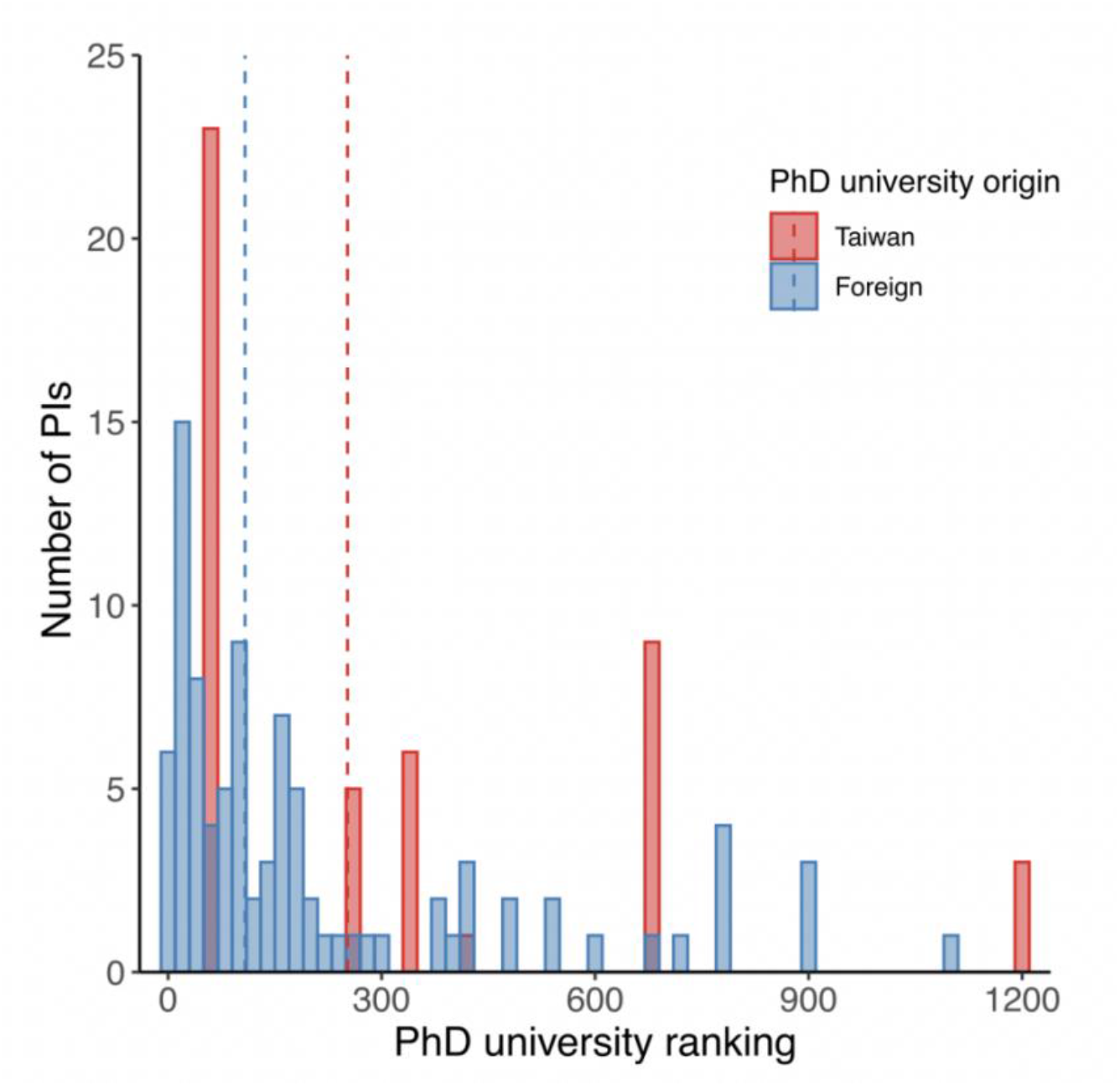
Distribution of the ranking of universities from which PIs obtained their PhD degrees. Dashed lines indicate medians of university ranking for Taiwanese (252) and foreign (108) PhD degrees

The academic performance before recruitment (“Before” h-index for recruitment) was higher for PIs who landed tenure-track positions more recently, whereas the performance for promotion to full professor (“Before” h-index for promotion) remained constant over years (Table 1, Fig. 3*a*–*b*). Although male PIs had on average higher performance than female PIs before recruitment, no such gender difference was found before promotion. PhD university origin and ranking had no effect on the performance either before recruitment or before promotion (Table 1).

**Table 1.**
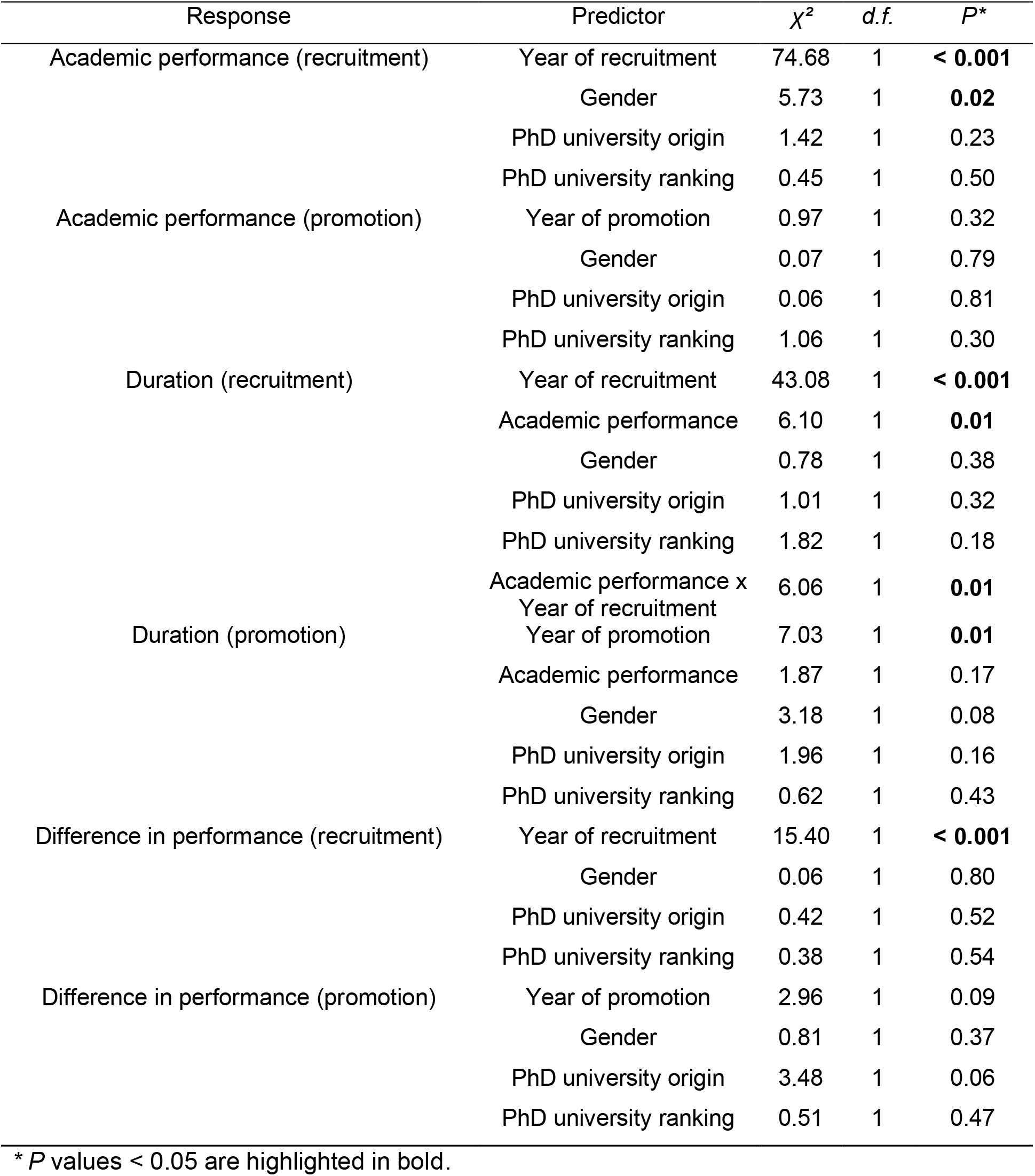
Results of the LMMs (type III sum of squares) on academic performance before recruitment/promotion (“Before” h-index), career duration before recruitment/promotion, and difference in performance before and after recruitment/promotion (“After” h-index — “Before” h-index)

**Fig. 3.**
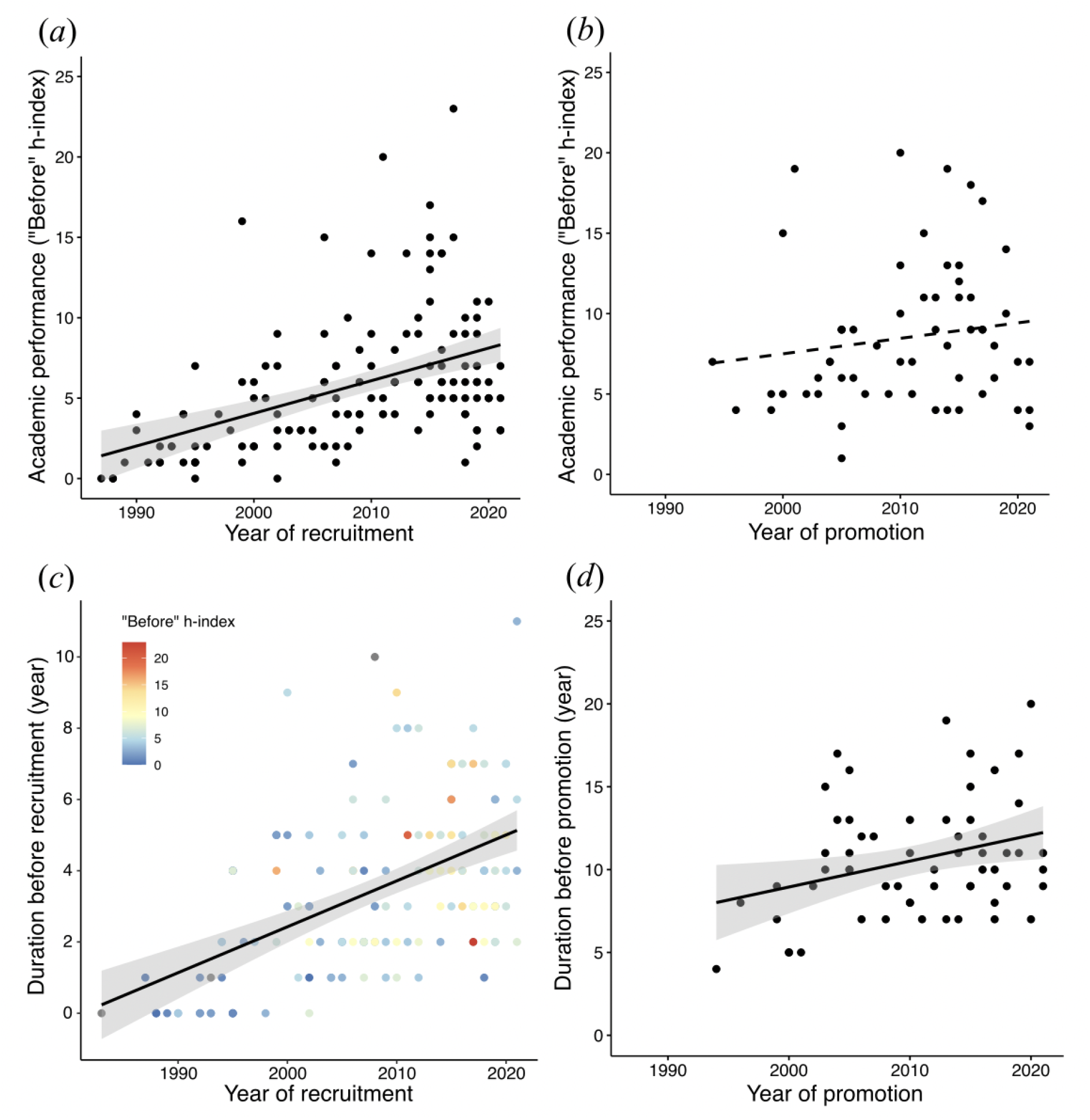
Temporal variations in academic performance (*a & b*) and career duration (*c & d*) before recruitment and promotion. Each point represents an individual PI, with points in (*c*) colored by “Before” h-index. Solid/dashed lines represent significant/non-significant relationships predicted from the LMMs; shaded areas indicate 95% confidence intervals

PIs who landed positions more recently spent more time post-PhD before recruitment, while higher academic performance reduced this duration (Table 1, Fig. 3*c*). On the other hand, PIs also spent more time before promotion to full professor in recent years, yet the duration was not related to the performance (Table 1, Fig. 3*d*). PhD university origin, ranking, and gender had no effect on the duration either before recruitment or before promotion (Table 1).

The difference in academic performance before and after recruitment (“After” h-index — “Before” h-index) decreased for PIs who landed positions more recently, while PhD university origin, ranking, and gender had no effect on the performance difference (Table 1, Fig. 4*a*–*b*). In contrast, the difference in performance before and after promotion to full professor was not associated with the year of promotion, PhD university ranking, or gender, yet the difference tended to be higher for PIs with foreign degrees compared to those with Taiwanese degrees (Table 1, Fig. 4*c*–*d*).

**Fig. 4.**
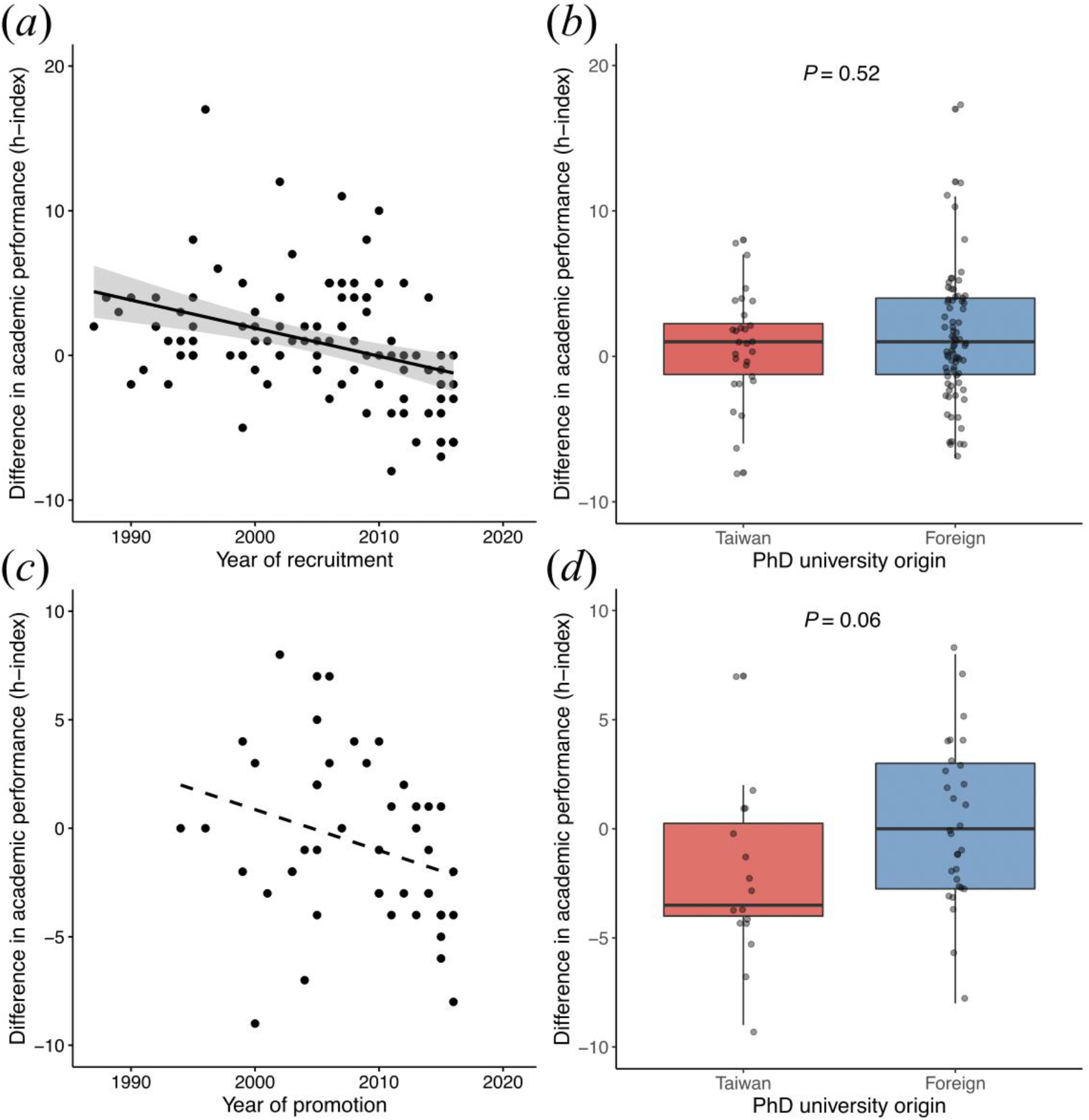
Difference in academic performance before and after recruitment (*a & b*) and promotion (*c & d*) (“After” h-index — “Before” h-index) in relation to the year of recruitment/promotion and PhD university origin. Each point represents an individual PI. Solid/dashed line represents significant/non-significant relationships predicted from the LMMs; shaded areas indicate 95% confidence intervals

## Discussion

Overall, we showed that the academic performance of PIs before recruitment as new faculty members increased over years, whereas the performance before promotion to full professor remained relatively unchanged. We also found that the duration both before recruitment and before promotion increased in recent years. These results provide empirical evidence supporting the suspicion that publication requirements and expectations have risen over time in the field of ecology and evolutionary biology in Taiwan, in line with many academic job markets worldwide (Rawat and Meena 2014; Warren 2019).

The increase in academic performance of PIs before recruitment suggests that the academic job market might have become increasingly competitive over time, which is likely driven by a relatively lower demand for tenure-track professors compared to the supply of new PhDs (Larson et al. 2014). Consequently, the duration post-PhD may be prolonged if the applicants are not competitive enough. However, higher academic performance could help shorten the time to land a position. Therefore, it would be important for early-career researchers to home in on publications in order to demonstrate their competence for academic success. In contrast, the performance of PIs before promotion to full professor remained similar over years, suggesting that the requirements for promotion might not have changed much over time. Interestingly, the time to full professor has lengthened in recent years but was not affected by academic performance, possibly due to increasing consideration of accomplishments such as teaching and administrative services by employment institutes in addition to research outputs. Such different patterns in academic performance and career duration between recruitment and promotion phase are likely due to applicants facing increasing competition with others during recruitment and thus higher performance would be advantageous for securing a position, whereas getting a promotion depends mainly on individual PI meeting the institutes’ requirements rather than comparing with others’ performance.

We found that the average performance of a new male PI was higher than that of a new female PI. This may result from higher standards for evaluating the suitability of a potential faculty member for males compared to females (Symonds et al. 2006). Alternatively, it could be due to employment institutes striving to recruit female applicants to enhance gender equity despite the likelihood of female applicants having a lower performance than their male competitors, which can be exacerbated by implicit bias and stereotype threats that females face in biological sciences (Salerno et al. 2019). However, the performance expectations for promotion to full professor did not differ between male and female PIs, indicating that after recruitment, especially when gender equality is enhanced, individual performance is the key to further promotion regardless of gender. Contrary to a previous study showing that researchers from higher-ranked institutes become PIs faster compared to those from lower-ranked institutes (van Dijk et al. 2014), we found no evidence of PhD university origin and ranking influencing the career duration either before recruitment or before promotion. Instead, our results suggest that academic performance during PhD and/or post-PhD period may be more important in determining the academic success compared with the prestige of education itself.

The difference in performance before and after recruitment decreased over years. Specifically, PIs in earlier years had on average higher h-indexes after recruitment than before recruitment, yet such a “performance boost” has declined in recent years. This could be due to increasing teaching and administrative demands of new PIs, reducing their time available for research. Surprisingly, we found that PIs performed consistently before and after recruitment regardless of their PhD university origin or ranking. However, PIs with domestic PhD degrees did show a decrease in performance after promotion to full professor compared to before promotion, whereas PIs with foreign PhD degrees had relatively consistent performance before and after promotion. One possible explanation is that the training and experiences from foreign universities may have equipped those PIs with greater professional abilities, which together with international connections and collaboration opportunities, help maintain their performance.

It is noteworthy that recruitment is a complicated process involving not only academic performance *per se* but also other considerations such as the suitability of applicants to the research areas of opening positions. Although our study showed increasing academic performance for recruitment over years, we do not intend to discourage the academic community with such results. Indeed, variations in h-index during recruitment phase indicate that it is still possible for an applicant with relatively low h-index to land a position. Moreover, besides research performance, other aspects of academic achievements, including teaching, mentoring, and social outreach, also constitute a significant part of a researcher’s career, and we stress that balancing these different aspects would be necessary for a more holistic professional development. Finally, our analyses were based on PIs in ecology and evolutionary biology. Since the nature of academic job markets can vary considerably among different fields of biology (Larson et al. 2014), the results should be interpreted carefully when applied to the fields outside the scope of this study. In conclusion, our findings confirm that succeeding in academia has become more challenging, with publication requirements and career duration both increasing over years. In the face of increasingly competitive academic job markets, boosting performance is a key to career success in academia.

## Acknowledgements

We thank Ming-Yang Megan Chang for constructive discussion and comments. S.-J.S. was supported by National Taiwan University and Ministry of Science and Technology, Taiwan (111WXA0310022).

## Statements and Declarations

### Competing interests

The authors declare no competing interests.

### Footnotes

Please note that this manuscript has also been posted on *bioRxiv* (Hsu et al. 2022) at https://www.biorxiv.org/content/10.1101/2022.01.31.478501v2, following the Springer Nature preprint sharing policy. It has also been added to the reference list.

### Funding

No funding was received for conducting this study.

### Authors’ contributions

G.-C.H. and S.-J.S. conceived the study; W.-J.L. and S.-J.S. collected the data; G.-C.H. and S.-J.S. analyzed the data. All authors were involved in writing the manuscript.

## Notes

### Competing Interest Statement

The authors have declared no competing interest.

### Summary of Updates

We have updated some of our results and analyses, as well as the discussion section.

## References

Acton, S.E., Bell, A.J., Toseland, C.P. & Twelvetrees, A. (2019). A survey of new PIs in the UK. eLife, 8.

Bates, D., Maechler, M., Bolker, B. & Walker, S. (2015). Fitting linear mixed-effects models using lme4. R package version.

Brischoux, F. & Angelier, F. (2015). Academia’s never-ending selection for productivity. Scientometrics, 103, 333–336.

Chen, N. (2021). “Why should a ‘foreigner’ be better than me?”: preferential practices in junior academic faculty recruitment among mainland Chinese universities. Tertiary Education and Management, 1–25.

Cyranoski, D., Gilbert, N., Ledford, H., Nayar, A. & Yahia, M. (2011). Education: The PhD factory. Nature, 472, 276–279.

Garfield, E. (1996). What Is The Primordial Reference For The Phrase “Publish Or Perish”? The Scientist, 10, 11.

Ghaffarzadegan, N., Hawley, J., Larson, R. & Xue, Y. (2015). A Note on PhD Population Growth in Biomedical Sciences. Systems Research and Behavioral Science, 32, 402–405.

Glänzel, W. (2006). On the h-index – A mathematical approach to a new measure of publication activity and citation impact. Scientometrics 2006 67:2, 67, 315–321.

Hirsch, J.E. (2005). An index to quantify an individual’s scientific research output. Proceedings of the National Academy of Sciences, 102, 16569–16572.

Hsu, G.-C., Lin, W.-J. & Sun, S.-J. Increased academic performance and prolonged career duration among Taiwanese academic faculty in ecology and evolutionary biology. bioRxiv, doi: https://doi.org/10.1101/2022.01.31.478501.

Larson, R.C., Ghaffarzadegan, N. & Xue, Y. (2014). Too many PhD graduates or too few academic job openings: The basic reproductive number R0 in academia. Systems Research and Behavioral Science, 31, 745–750.

Laurance, W.F., Useche, D.C., Laurance, S.G. & Bradshaw, C.J.A. (2013). Predicting Publication Success for Biologists. BioScience, 63, 817–823.

Lenth, R. v. (2021). emmeans: Estimated marginal means, aka least-squares means. R package version 1.7.1. R Foundation for Statistical Computing.

National Science Foundation. (2018). Science and Engineering Indicators. NSB-2018-1. Available at: https://www.nsf.gov/statistics/seind/. Last accessed 6 February 2022.

Pauly, D. & Stergiou, K.I. (2005). Equivalence of results from two citation analyses: Thomson ISI’s Citation Index and Google’s Scholar service. undefined, 5, 33–35.

R Development Core Team. (2014). R: A language and environment for statistical computing. R Foundation for Statistical Computing.

Rawat, S. & Meena, S. (2014). Publish or perish: Where are we heading? Journal of Research in Medical Sciences : The Official Journal of Isfahan University of Medical Sciences, 19, 87.

Ryan Haley, M. (2012). Rank variability of the Publish or Perish metrics for economics and finance journals. http://dx.doi.org/10.1080/13504851.2012.697115, 20, 830–836.

Schillebeeckx, M., Maricque, B. & Lewis, C. (2013). The missing piece to changing the university culture. Nature Biotechnology 2013 31:10, 31, 938–941.

Shin, J.C. & Kehm, B.M. (2013). Institutionalization of world-class university in global competition. Institutionalization of World-Class University in Global Competition, 1–301.

Swihart, R.K., Sundaram, M., Höök, T.O. & Dewoody, J.A. (2016). Factors affecting scholarly performance by wildlife and fisheries faculty. The Journal of Wildlife Management, 80, 563–572.

Symonds, M.R.E., Gemmell, N.J., Braisher, T.L., Gorringe, K.L. & Elgar, M.A. (2006). Gender Differences in Publication Output: Towards an Unbiased Metric of Research Performance. PLOS ONE, 1, e127.

van Dijk, D., Manor, O. & Carey, L.B. (2014). Publication metrics and success on the academic job market. Current biology : CB, 24.

Warren, J.R. (2019). How much do you have to publish to get a job in a top sociology department? Or to get tenure? Trends over a generation. Sociological Science, 6, 172–196.

Witteman, H.O., Hendricks, M., Straus, S. & Tannenbaum, C. (2019). Are gender gaps due to evaluations of the applicant or the science? A natural experiment at a national funding agency. The Lancet, 393, 531–540.

Xue, Y. & Larson, R.C. (2015). STEM crisis or STEM surplus? Yes and yes. Monthly labor review, 2015.

